# Diverging Hydroclimatic Trends in Global Tropical Dryland Ecosystems Based on ERA-5 and CHIRPS Analysis Data

**DOI:** 10.64898/2026.06.23.734075

**Authors:** Arturo Sanchez-Azofeifa, Kayla Stan, Hendrik F. Hamann

## Abstract

Tropical dryland ecosystems are highly biodiverse and fragmented and are experiencing significant anthropogenic and climatic changes. With increasing extremes in temperature and precipitation, coupled with significant alteration, these ecosystems are at greater risk of increased exposure and vulnerability to climatic change; however, little work has quantified the climatic shifts occurring within these ecosystems globally. Here, we aim to fill this gap by using the ERA-5 reanalysis and CHIRPS precipitation data to quantify changes in essential climatic variables in tropical drylands since 2000. Overall, we find that regional pressures differ, with tropical dry forests, savannas, and shrublands becoming hotter and drier in the Neotropics and parts of the Afrotropics and Australasia. By contrast, the tropical dry forests in the Indomalayan, Oceania, and Nearctic are experiencing hotter and wetter conditions. Globally, though, these ecosystems are experiencing more change than the global average, suggesting they may be approaching tipping points in their resilience, ultimately shrinking the area where they can survive.

## Introduction

Tropical dryland ecosystems have experienced some of the highest levels of anthropogenic alteration in the last two decades and remain among the most threatened ecosystems (Portillo-Quintero et al. 2010; Schröder et al. 2021; Stan et al. 2024). These biomes provide many ecosystem services, including erosion control, carbon sequestration, climate and water regulation, and high biodiversity; however, their favourable climatic conditions make them attractive for both population growth and agriculture (Andrade et al. 2020; Calvo-Rodriguez et al. 2016; Nelson et al. 2020). In fact, these biomes are located adjacent to highly populated areas and are consequently subject to stress from anthropogenic alterations, agricultural encroachment, and overgrazing (Miles et al. 2006; Portillo-Quintero et al. 2010; Siyum 2020). This alteration has fragmented many of these sensitive ecosystems, with less than ⅓ of the original tropical dry forests remaining and less than 20% of the remaining forests located in non-fragmented areas (Stan et al. 2024). Beyond substantial anthropogenic stressors, these ecosystems have also experienced shifts in their hydroclimatic regimes, with increased forest mortality and reduced resilience attributed to increased droughts and disturbances such as fires and insect outbreaks (Hartung et al. 2021; Siyum 2020; Suresh et al. 2010).

Recent studies indicate that dryland ecosystems worldwide may experience increased aridity under climate change, while others suggest that plant functional types will shift into other ecosystems under these new climatic regimes (Kim et al. 2023; Moura et al. 2023; Park et al. 2018). However, at the regional scale, there is a discrepancy between the current and projected resilience of tropical dry forests to climate change. Within the Neotropics, some sites are resilient to climate change and have experienced increases in growing-season length and overall productivity (Stan et al., 2019; Stan et al., 2021). These mixed trends of some regions experiencing increases in productivity and growing season length, while other regions are experiencing decreased growing seasons, are also found on a global scale (Guo et al. 2022; Xu et al. 2023). Other studies suggest that tropical dry forests, along with many other forested ecosystems, are nearing their climatic limits, particularly concerning temperature and precipitation (Doughty et al. 2023).

This combined impact of anthropogenic alteration and fast-approaching climatic thresholds of resilience may indicate that these dry ecosystems are nearing a tipping point beyond which they may no longer be resilient (Abdaki et al., 2025). Despite the importance of these ecosystems and the extent of the globe they cover, large scale models typically focus only on temperature or precipitation changes, despite local-scale work indicating that dry ecosystems are sensitive to belowground characteristics, vapour pressure deficit, and light (Rankine et al. 2024; Werden et al. 2022; Yang et al. 2023). There remains a gap in research in characterizing large-scale climatic changes in tropical dry ecosystems using reanalysis data and on understanding regional differences in these changes.

Here, we aim to fill this gap by using state-of-the-art reanalysis data from ERA-5 to characterize changes in the essential climatic variables in global tropical dry ecosystems. Essential Climate Variables provide an integrative, holistic view of how the climate is changing in these ecosystems, and ERA-5 enables us to calculate changes in tropical dry climates over the past two decades. Overall, and perhaps not surprisingly, we found that tropical dry ecosystems across the globe do not experience consistent climatic pressures, with some sites showing a trend towards aridification. In contrast, others show increases in precipitation and humidity. While these selective pressures are different, ultimately, both are likely to push the tropical dry forests towards their limits of resilience and reduce the area that is optimal for this unique and highly biodiverse ecosystem.

## Data and Methods

### Study Area

Tropical dryland ecosystems include dry forests, shrublands, and savannas (Pennington et al. 2018; Stan et al. 2024). Broadly speaking, however, these ecosystems occur in year-round warm climates, with average temperatures greater than 25°C and an extended dry season with less than 100 mm of precipitation per month for 3-8 months (Portillo-Quintero 2010; Ocon et al. 2020). Savanna biomes are typically characterized by an open canopy with <80% canopy cover and a grassy understory, whereas tropical dry forests typically have a closed canopy (Pennington et al. 2018). Dry forests contain a mix of deciduous and evergreen trees; however, the vegetation is typically dominated by deciduous tree species. A universal definition of dry forests is lacking, and their high seasonality, paired with high cloud cover during productive periods, often leads to misclassification as savanna regions because most mapping and monitoring occur during the dry season. These discrepancies are particularly noticeable between Africa and Asia, where similar woodland structures are classified as savannas and forests, respectively (Pennington et al. 2018).

Given the contentious and often inconsistent distinctions between tropical dry forests and tropical dry savannas, this study aims to provide a broader definition of the impact of climate change on tropical dry ecosystems. To do this, the RESOLVE 2017 ecoregions dataset was used, with the Tropical and Subtropical Dry Broadleaf Forests (TSDBF) and Tropical and Subtropical Grasslands, Savannas, and Shrublands (TSGSS) ecoregions specifically targeted (Dinerstein et al. 2017). These two biomes cover 114 unique ecoregions spanning 105 countries. The two ecoregions were considered separately to assess differences between the biome types. The RESOLVE dataset is an updated version of the Terrestrial Ecoregions of the World, as defined by Olson et al. (2001). The ecoregions are also subdivided into six macroecological locations: Neotropical, Afrotropic, Indomalayan, Australasia, and Nearctic.

### Climate Data

Essential Climate Variables (ECVs) are variables defined as necessary to monitor and characterize Earth’s climate more holistically (Bojinski et al., 2014). These variables are essential for predicting future climates and guiding future climate adaptation and mitigation strategies. These variables are defined by the Global Climate Observing System (GCOS) and currently include 55 variables that span the atmospheric, marine, anthropogenic, and terrestrial spheres. The Copernicus Climate Change Service at the European Center for Medium-Range Weather Forecasts has created a state-of-the-art dataset using its ERA-5 reanalysis data archive (Hersbach et al., 2020). This continuous, global, hourly dataset of observations begins in 1979 (with a backcast extending to 1940) at a 0.25° resolution and is generated by assimilating remote-sensing datasets into the model (∼30 km resolution).

In this study, we used seven global climate variables from the ERA-5 archive. The ECVs used in this study included surface temperature (T; °C), dew point (DP;°C), solar radiation (SR; J/m²/hr), cloud cover (CC), atmospheric water content (AWC; kg/m²), atmospheric water vapor content (AWVC; kg/m²), and volumetric soil water content (VSWC; m_3_/m_3_). These ECVs were selected because they are most commonly used to monitor climate in drylands and provide an intuitive understanding of past, present, and future climatic changes. Precipitation data were collected from the Climate Hazards Infrared Precipitation with Stations (CHIRPS) dataset. CHIRPS includes precipitation data from 50 °N to 50 °S, spanning 38 years at a 0.05 ° resolution (Funk et al., 2015). All climate variables were collected from 1997 to 2020 between 50 °N and 50 °S for consistency in the analysis parameters.

While compiled and reanalyzed data, such as ERA-5 and CHIRPS, have biases at a local scale, these continuous time series across a single dataset allow us to assess climate variable trajectories in ecosystems and compare these trends over time and across regions (Bonshoms et al. 2022; Dommo et al. 2022; Guzmán et al. 2024). These comprehensive, state-of-the-art products enable assessment of trends over long timescales and across the globe.

### Temporal Trends & Statistical Analysis

The hourly data were assessed pixel by pixel, and annual changes in each variable were calculated using linear regression. To calculate the regression coefficients, we utilized IBM PAIRS Geoscope. IBM PAIRS enables user-defined functions, such as linear regression, to be applied to ERA-5 and CHIRP data without downloading any raw data (Lu et al., 2016). Instead, the regression coefficients and uncertainties for each pixel were returned directly. During the regression analysis, IBM PAIRS processed ∼13TB of ERA-5 and CHIRPS data in less than 3 hours, yielding, for each pixel, the regression coefficients (slope, intercept, and standard deviation).

The regression coefficients were then subset into Subtropical and Tropical Broadleaf Dry Forests (TSBDF), Tropical and Subtropical Grasslands, Savannas and Shrublands (TSGSS), and all other land masses. The data were further subset by region and included the Neotropical, Afrotropic, Indomalayan, Australasia, and Nearctic realms. The Paleoarctic, the world’s largest biogeographic region, does not contain TSBDF or TSGSS and was therefore excluded from the analysis. The subsets were based on ecoregion extents from the Ecoregion datasets and were not filtered by land-cover type to ensure consistency. The regression coefficient distributions of each data subset were compared using the Kruskal-Wallis test and Post-hoc Dunn test (MacFarland & Yates, 2016). The regression coefficients were not normally distributed (p<0.05); therefore, parametric statistics such as the Bayes t-test or ANOVA were not used.

Finally, to understand the collective change in the climatic niches occupied by the tropical dry ecosystems across realms, a principal component analysis (PCA) was performed on the regression coefficients. All variables were tested for fitness for running dimension reduction processes using the Kaiser-Meyer-Olkin test and the Bartlett’s Test of Sphericity with the psych package in R. The average KMO of all factors was 0.71, with all variables having KMO values greater than 0.5, while the Bartlett p-value was <0.01, thus indicating that the data is suitable for a PCA. The PCA was performed on the data in R using the FactoMineR package (Husson et al., 2024). The total explanatory power of each component was assessed, and the distribution of each biome and realm on these new dimensions was compared to determine the collective climate trends within each biome and across realms.

## Results

Over the past four decades, trends in key hydrologically relevant climatic variables, such as solar radiation, cloud cover, dew point, and atmospheric water content, have diverged (Figure 1). Sub-Saharan Africa, Central America, India, and much of Southeast Asia have experienced increases in cloud cover and decreases in solar radiation, with some areas seeing decreases of up to 5700 J/m^2^/yr. In contrast, much of South America, south-central Africa, and much of Australia experienced opposite trends, with increases in solar radiation of up to 4300 J/m^2^/yr. These diverging trends indicate that similar biomes across the globe may face different pressures and be required to adapt to changing climatic conditions.

**Fig 1.**
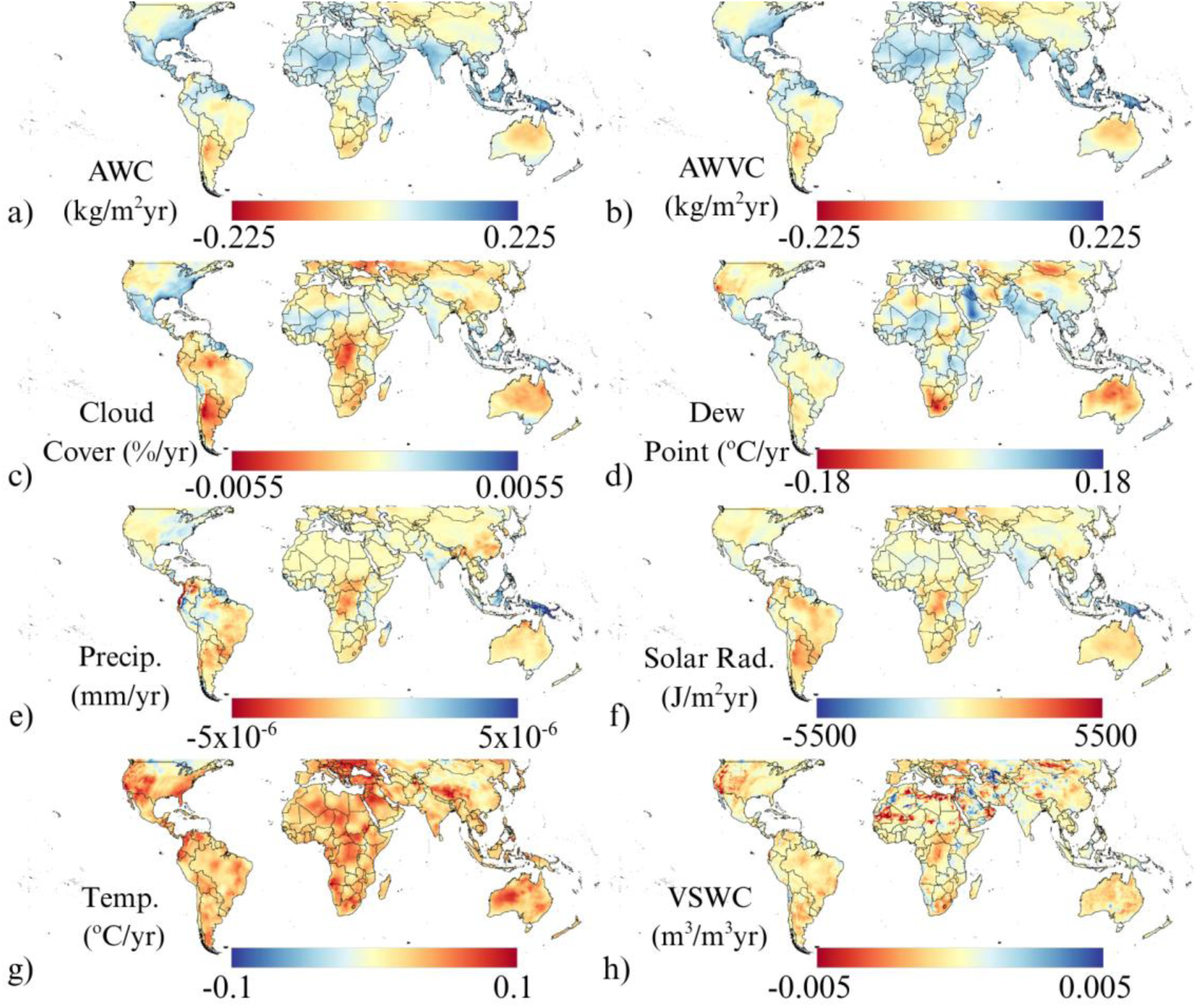
Essential climatic annual trends on landmasses around the world. The trends encompass annual rate of change in a) atmospheric water content, b) atmospheric water vapour content, c) cloud cover, d) dew point, e) precipitation, f) solar radiation, g) temperature, and h) volumetric soil water content from 2000 to 2019. The trends were estimated by regressing the change over time between 50° and −50° latitudes using a 0.13-degree grid.

### Global Comparison of Tropical Dry Biomes Climatic Trends

When we consider global trends in ECVs across TSBDF and TSGSS, we observe annual increases of 0.027 °C/yr and 0.03 °C/yr, respectively (Figure 2, Table 1). There were also consistent increases in atmospheric water content: 0.074 kg/m²/yr for TSDBF and 0.0009 kg/m²/yr for TSGSS. The increase in atmospheric water vapour content mirrors this trend. Although both biomes experienced warming, along with the rest of the landmasses, divergent patterns were observed between the TSBDF and TSGSS in changes in radiative forcing and other essential climatic variables. These differences were evident when comparing these biomes with other central-latitude land areas.

**Fig 2.**
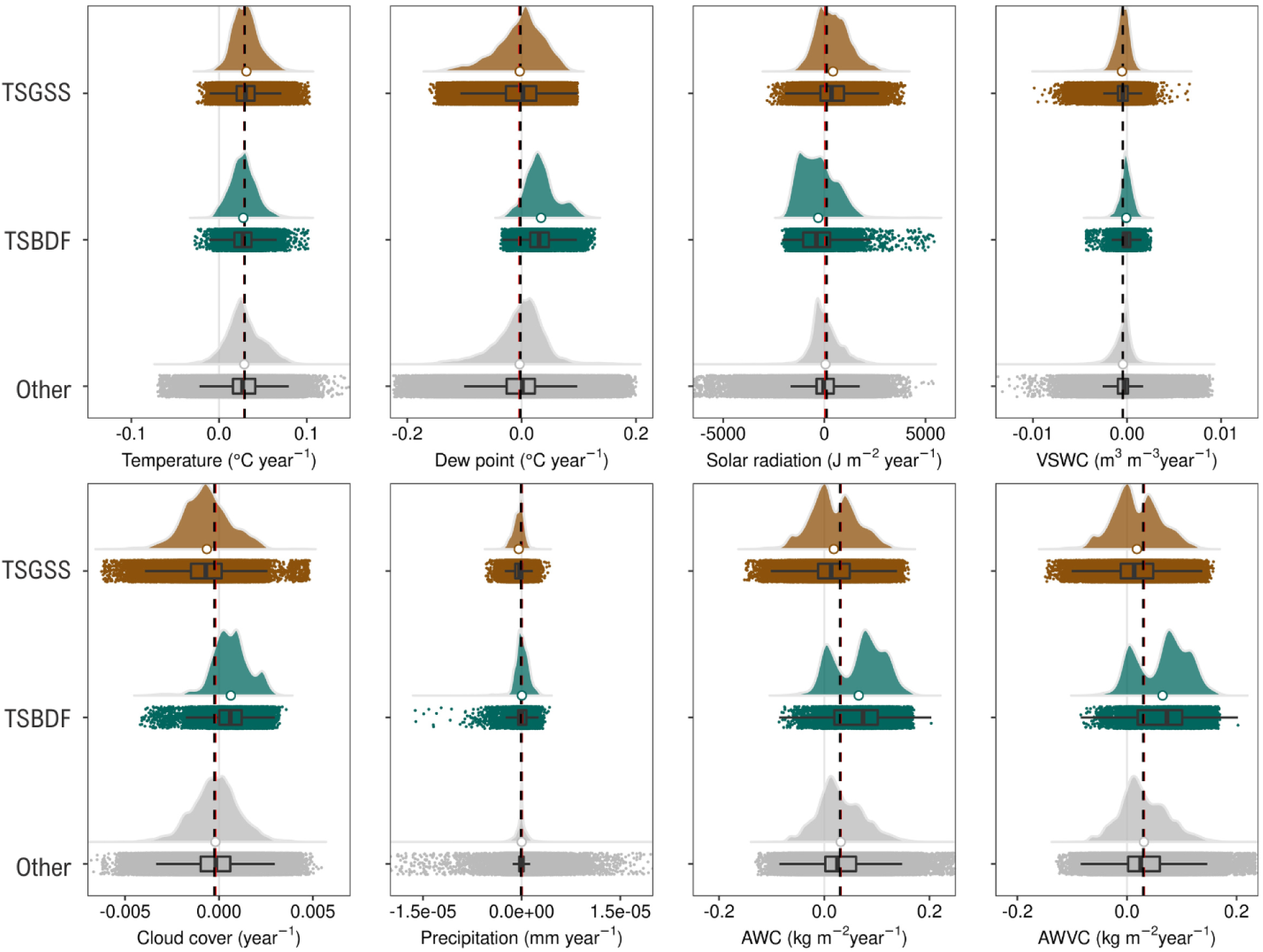
Raincloud plot comparing ECV trends across dry biomes and other landmasses. The black and red dashed lines represent the averages of global trends and landmasses, excluding dry biomes. Green represents TSBDF, brown represents TSGSS, and gray represents other land masses without TSBDF and TSGSS.

**Table 1:** Comparison of changes in essential climatic variables within tropical dry biomes and other land masses.

| Essential Climatic Variable | $\Delta$ Other | $\Delta$ TSBDF | $\Delta$ TSGSS |
| --- | --- | --- | --- |
| Atmospheric Water Vapour Content<br>(kg/m<sup>2</sup>/yr) | +0.036 | +0.074 | +0.009 |
| Atmospheric Water Content<br>(kg/m<sup>2</sup>/yr) | +0.036 | +0.075 | +0.009 |
| Cloud Cover (%/yr) | $2.22 \times 10^{-5}$ | $4.80 \times 10^{-4}$ | $-6.88 \times 10^{-4}$ |
| Dew Point (°C/yr) | +0.014 | +0.029 | +0.001 |
| Precipitation (mm/yr) | $+5.3 \times 10^{-8}$ | $+7.1 \times 10^{-8}$ | $-3.4 \times 10^{-7}$ |
| Solar Radiation (J/m<sup>2</sup>/yr) | -119.2 | -320.9 | +388.3 |
| Temperature (°C/yr) | +0.026 | +0.027 | +0.030 |
| Volumetric Soil Water Content<br>(m<sup>3</sup>/m<sup>3</sup>/yr) | -0.0002 | 0.0000 | -0.0005 |

In the TSBDF, changes in ECVs were similar in direction, though often exceeded the average change relative to the larger global mean changes during the same period. These changes include increases in atmospheric water content and cloud cover and decreases in solar radiation. The TSBDF experienced the largest increase in Atmospheric Water Vapour Content (AWVC; +0.075 kg/m²/yr), whereas the global average increase was only +0.036 kg/m²/yr. Cloud cover increased by 0.0005 %/yr, which is 25 times higher than the global mean change (+0.00002 %/yr). This increase in cloud cover is paired with a twofold decrease in solar radiation relative to the global average of −321 J/m²/yr. However, these changes do not affect precipitation or soil water content in the TSBDF and only cause small drops in global soil water content.

The TSGSS regions deviated strongly from the global and TSBDF trends in terms of directionality and magnitude. While TSGSS’s average warming rate (+0.039°C/yr) exceeded both the global and TSBDF trends, it experienced the lowest increase in atmospheric water content (0.009 kg/m²/yr). Additionally, the TSGSS showed gains in solar radiation (+388.3 J/m²/yr), decreases in cloud cover (−0.0006/yr), and the largest decrease in soil water content (−0.0005 m^3^/m^3^ /yr). These values differ widely from the global mean solar radiation decline of –119.2 J/m²/yr and the –320.9 J/m²/yr decline within TSBDF.

### Regional Variations in Tropical Dry Biomes

While there is a divergence in climatic stress globally between the TSBDF and TSGSS, this difference is even greater at the regional level. Additionally, climatic stress varies significantly across biomes, indicating distinct stresses and vulnerabilities faced by ecoregions within each biome.

Within the TSBDF biome, the Nearctic realm showed the strongest increases across most ECVs considered, and all of these increases were significantly different from those in other regions (p<0.001). The temperature increase was the highest within the TSBDF and across the other biomes (0.06°C/yr, p<0.001). The Nearctic dry forest also had the largest increases in atmospheric water content (+0.13 kg/m²/yr), atmospheric vapour water content(+0.13 kg/m²/yr), cloud cover (+0.002%/yr), and dew point (+0.08 °C/yr). These increases exceed the maximum regional changes in the TSGSS and other landmasses, except for cloud cover. Although not the most extreme, these increases in the Nearctic are accompanied by little change in precipitation (2×10^-7^ mm/yr) and by decreases in solar radiation (−969 J/m²/yr). Therefore, warming in the Nearctic TSBDF is accompanied by increased humidity, but this has not translated into more water on the ground.

In contrast, the Australasian realm TSBDF experienced the largest increase in solar radiation (+219 J/m²/yr), which was paired with decreases in cloud cover (−0.0001/yr) and small but significant decreases in both precipitation (−2×10^-7^ mm/yr) and soil water content (−2×10^-5^ m^3^/m^3^/yr). This pattern of increased radiation and decreased cloud cover is also observed in the Afrotropics (+49 J/m²/yr and −0.0003/yr, respectively), although it is less pronounced than the trend in Australasia. However, the Afrotropics did not experience a corresponding decrease in precipitation (+2×10^-7^ mm/yr) or in soil water content (+3×10^-4^ m3/m^3^ /yr). While Afrotropical dry forests have experienced increases in radiative forcing, this has not translated into reduced water availability. Given the ongoing trends in Australasian dry forests, which have both increased radiative forcing and decreased water availability, there may be concern that this trend will occur in the Afrotropics in the future.

**Fig 3.**
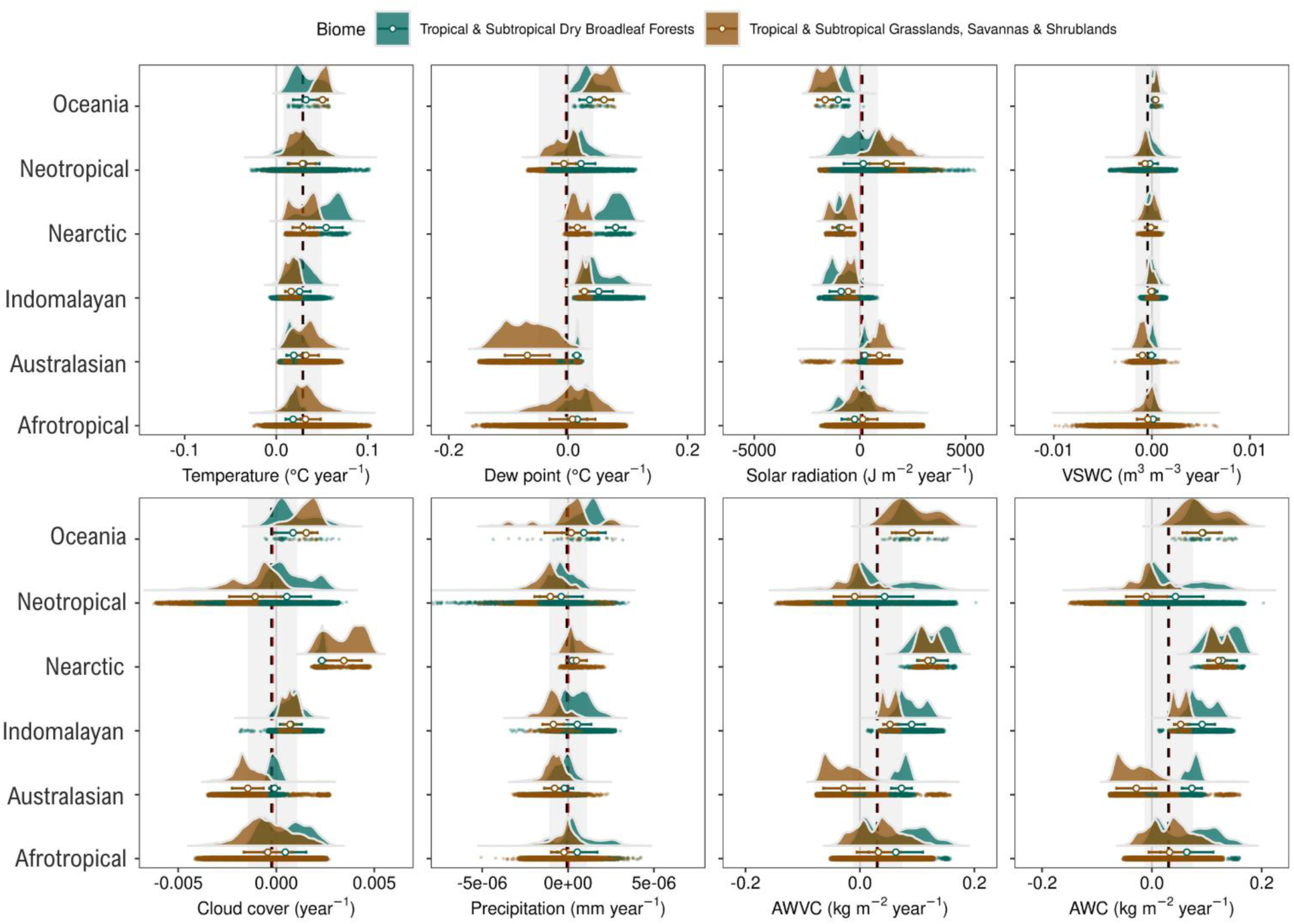
Raincloud plot comparing ECV trends across dry biomes in different realms and other landmasses. Black and red dashed lines represent the average of global trends and landmasses, respectively, excluding dry biomes.

The Indomalayan and Oceanian TSBDF showed mixed patterns, with both regions experiencing moderate increases in temperature (+0.03 °C/yr) and strong declines in solar radiation (Indomalayan: –886 J/m²/yr; Oceania: –1,036 J/m²/yr). These realms also experienced small increases in soil water content (Indomalayan: 6 x 10^-5^ m^3^/m^3^ /yr, Oceania: 2 x 10^-4^ m^3^/m^3^/yr), and the largest increases in precipitation (Indomalayan: +5 x 10^-7^ mm/yr, Oceania: 1 x 10^-6^ mm/yr). A significant increase in cloud cover accompanies increased precipitation and decreased solar radiation (Indomalayan: 6 x 10^-4^/yr; Oceania: 0.001/yr; p<0.001). These trends may indicate changes in the monsoon season and in precipitation during the wet season in both regions.

When we consider the TSGSS biome, trends differ, particularly in changes in water availability. Most regions experienced decreased precipitation and soil water content. The Australasian TSGSS showed the greatest variation, being the only region to experience decreases in atmospheric volumetric water content (−0.02 kg/m²/yr) and dew point (−0.05 °C/yr). These decreases are also accompanied by a drop in precipitation (−7.1 x 10^-7^ mm/yr), the largest regional decreases in cloud cover (−0.001/yr) and soil water content (−9 x 10^-4^ m^3^/m^3^ /yr). Only solar radiation and temperatures have risen in the region (SR: +925 J/m²/yr; T: +0.027 °C/yr), indicating a warming and drying trend in this water-limited biome.

The Neotropical TSGSS also experienced large increases in both solar radiation (+853 J/m²/yr) and temperature(+0.028 °C/yr). The water availability in the vadose zone was also reduced, with decreases in soil water content (−5.4×10^-4^ m^3^/m^3^ /yr) and the largest decrease in precipitation (−8.1×10^-7^ mm/yr). These trends in water content do not match the atmospheric water vapour content (0.007 kg/m²/yr), which has moderately increased in the Neotropical TSGSS over the last four decades. These conditions indicate increased evapotranspiration, which is not replenished by precipitation within the biome.

The Nearctic and Oceania TSGSS both showed opposite changes in climatic conditions, with the Nearctic experiencing the greatest increases in cloud cover (+0.004 /yr) and precipitation (+3.4 x 10^-7^ mm/yr). Oceania had the greatest increases in temperature (+0.05 °C/yr) and soil water content (1.1 x 10-4 m3/m^3^/yr), paired with the largest decrease in solar radiation (−1,339 J/m²/yr). The increases between the Nearctic TSGSS and Oceania TSGSS in cloud cover, atmospheric volumetric water content, precipitation, and soil water content, along with the decrease in solar radiation, were not significantly different (p > 0.13). The Indomalayan TSGSS also exhibited similar trends, with no significant difference between this realm and the Nearctic and Oceania for both changes in solar radiation and soil water content (p > 0.39). The Indomalayan TSGSS was also not significantly different from the Oceania TSGSS in terms of increases in atmospheric volumetric water content, cloud cover, or dew point (p > 0.20). Overall, these realms have warmer, wetter, and more humid conditions, with the Nearctic TSGSS experiencing the largest change, followed by the Oceania and Indomalayan TSGSS.

Overall, the changes to the regional TSBDF ECVs were more isolated, with 12% of changes not significantly different (p > 0.05). There was more overlap in the TSGSS changes, particularly between the Indomalayan, Oceania, and Nearctic realms. In these biomes, 19% of the changes between regions were not significantly different (p > 0.05). Taken together, this does indicate a high level of regionally specific changes within the biomes, which may indicate divergent climatic pressures on these ecosystems.

### Clustering of Climatic Spaces

Principal Component Analysis (PCA) showed strong overlap between changes in TSGSS and TSBDF; however, distinct clustering patterns were observed. Beyond this, there is a strong directional bias that clusters tropical dry ecosystems that do not occur in other biomes. The first three principal components accounted for 81.3% of the total variance, with the first, second, and third components explaining 45.2%, 23.8%, and 12.2%, respectively. In general, cloud cover, solar radiation, and precipitation were loaded onto the first component, with a corresponding increase in cloud cover and precipitation. Changes in temperature, atmospheric water vapour content, and volumetric solid water content are loaded onto the second component, with a negative correlation between temperature and soil water content.

**Fig 4.**
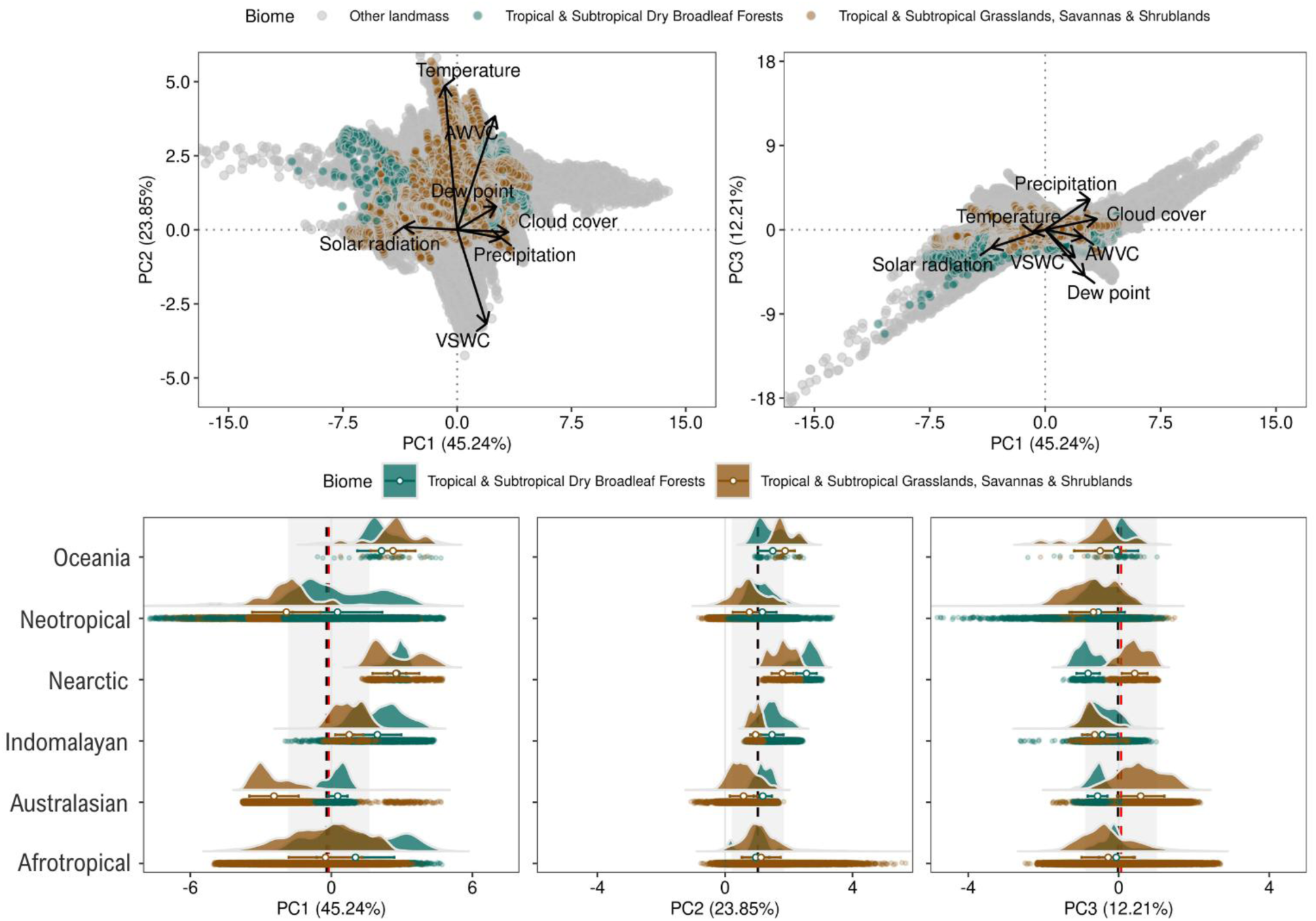
Variability in ECV trends across dry biomes in different realms and other land masses. PCA biplot with ECV plotted using the first two principal components (a) and the first and third components (b). PCA scores from the first (c), second (d), and third (e) components were compared between dry biomes using raincloud plots.

Generally, the TSBDF loads positively on the first component, with all regional averages also positive. However, there is substantial variability across the Neotropics, with some areas loading strongly in the negative part of the first component. The TSGSS was more variable in the first component, with Australasia and the Neotropics loading onto the negative side, indicating reduced cloud cover and precipitation and increased solar radiation. In contrast, the Indomalayan, Nearctic, and Oceania TSGSS clusters were on the positive side of the first component, indicating increased solar radiation. The Afrotropics showed the greatest variation in first-component values.

PCA’s second component is highly skewed in the positive direction for both the TSBDF and TSGSS. The TSGSS again shows greater variation but generally has slightly lower average positive values across realms than the TSBDF. Only the Oceania TSGSS had higher average values for this component, while the Afrotropics had similar averages but a much wider distribution. The TSGSS skews to the highest loadings on component two, but otherwise, neither the TSGSS nor the TSBDF has the largest outlier changes in the PCA. This is particularly true for the positive side of the first component and the negative side of the second component.

Taken together, the PCA shows that TSGSS shift to hotter, sunnier, and drier conditions, particularly in parts of the Afrotropical, Australasian, and Neotropical ecosystems, whereas many TSBDF ecosystems experience cloudier, hotter, and more humid conditions, particularly in the Nearctic, Indomalayan, and Oceania regions.

## Discussion

Our study found different climatic pressures not only between the TSGSS and TSBDF biomes, but also across realms. This contrasts with many future climate projections, which suggest that dryland areas will become hotter and drier, and that tropical dry ecosystems will experience greater drought stress (Kim et al. 2023; Park et al. 2018). While this may be true for the savanna and shrubland ecosystems of the Afrotropical, Neotropical, and Australasian realms, it is not true for the dry forest ecosystems in the Nearctic, Indomalayan, and Oceania regions. These results are consistent with recent work suggesting a mix of extended and shortened growing seasons across dry forests (Guo et al. 2022; Xu et al. 2023). Increases in precipitation, however, may not result in a more resilient tropical dry ecosystem.

Ecosystem responses to drying conditions have been studied more frequently in tropical dry forests, as models have suggested continued aridification in water-limited ecosystems. There is growing concern that water-limited ecosystems are close to their climatic limit in terms of aridity; however, long-term selective pressures suggest that some species in those ecosystems are well adapted to increased drought (Gonzalez-M et al. 2020). Greater investment in root systems or in expensive wood tissues related to hydraulic efficiency has been found to improve drought resilience and biomass dominance in these ecosystems and is common among species in this biome (Gonzalez-M et al. 2020; Werden et al. 2022). While droughts have been found to impact ecosystem productivity locally and regionally, these ecosystems typically recover from severe droughts within 1-2 years (Castro et al. 2018; Jiao et al. 2021; Stan et al. 2021; Yu et al. 2017). Some studies have suggested that multi-decadal droughts are required to push these ecosystems past their tipping point (Mendivelso et al. 2014). However, the resilience of dry forests to drought may be moderated by site factors, with drier sites taking longer to recover after severe drought and human disturbance (Hollunder et al. 2022; Pagotto et al. 2025, Zou et al. 2020). Our work suggests that these conditions may become increasingly concerning in some Australasian, Afrotropical, and Neotropical ecosystems. In regions with increased hot-drought conditions, species richness will likely decrease, and site homogeneity will increase, particularly at sites that have not invested as heavily in drought-tolerant strategies. They may also experience increased mortality and decreased above-ground biomass (Gonzalez-M et al., 2020; Manrique-Ascenio et al., 2024). This decrease in diversity is problematic because it will continue to degrade ecosystem functionality and reduce resilience (Hong et al. 2021). These patterns may also be exacerbated by increased forest disturbances, including fires and agricultural expansion, which have been found to increase in drier years (De Marzo et al., 2021).

While not all regions have experienced hotter, drier conditions, changes towards increasing precipitation do not necessarily indicate an increase in resilience for tropical dryland species. Increases in humidity and precipitation indicate either a shortening dry season or increased precipitation during the monsoon season. Models suggest that increases in precipitation in dryland ecosystems do not necessarily lead to increases in biomass and leaf area index, as these wetter sites are often limited by light and nutrient availability (Zhang et al. 2022). If the increased precipitation is anomalous and occurs during the monsoon season, it could also lead to structural overshoot that year,, depleting soil moisture and increasing drought stress and mortality in future years (Zhang et al., 2022). A shortening of the dry season may also reduce species richness by decreasing the competitiveness of drought-tolerant species and allowing the infiltration of species more commonly found in moist forests. Similar transitions have been studied with shrubland encroachment into grassland regions, but remainargely unstudied inathe tropical dry forest-w-t forest interface (Walter et al. 2,024). This type of infiltration is particularly of concern during early successional stages, where drought tolerance traits are more important, and the successional pathway difference between wet and dry forests is more pronounced (Poorter et al. 2021). Such a reduction in resilience has been found in the Indomalayan dry forests,, which share phenological and floristic characteristics with species found in the Neotropical dry forests (Kurten et al. 2017). The dominant tree species in this region require a substantially dry season with <50 mm of rain for at least four months (Deb et al. 2017; Kurten et al. 2017), and increased precipitation makes them less resilient and competitive. With increased precipitation in the Indomalayan region, the suitable habitat for these key species is currently under threat.

Increased precipitation may also be linked to more extreme precipitation events, which could result in flooding and damage to species not accustomed to high water influxes, similar to their Indomalayan aseasonal forest counterparts (O’Brien et al. 2024). There is, however, limited research on the impact of flooding or increased precipitation on tropical dryland ecosystems. In temperate deciduous forests, increased precipitation has been found to lead to early dormancy (Xie et al., 2015). In savannas, above-average rainfall is associated with reduced productivity, and in grasslands, extremely wet years decrease species richness (Kanniah et al. 2013; Perez et al. 2024). However, the maximum hydric limit in tropical dryland forests and shrublands remains poorly understood. This limits our ability to predict future tropical dryland resilience under changing climatic conditions, including regional increases in water availability and drought (Knapp et al., 2017; Wang and Collins, 2024).

Overall, many tropical dryland ecosystems have experienced significant changes in their climatic conditions over the past few decades. These transitions, if they continue, are pushing forests toward tipping points where they are no longer resilient, with drier ecosystems potentially experiencing increased mortality and shrubification. At the same time, wetter sites may be at risk of invasion by wet forest species. These ongoing climatic changes will likely increase pressure on ecosystem functions, with potential consequences for flora, fauna, and human economic activities that rely on these sensitive ecosystems. Generalizations of the pressures that ecosystems face over biomes or globally reduce our ability to predict what will happen in the future. Our work shows that significant and distinct changes occur across tropical dryland biomes, and these climatic pressures are even more nuanced when reduced to individual ecoregions. Moving forward, regionally and locally based research is necessary to understand ecosystem resilience to factors beyond temperature increases and precipitation reductions.

Additionally, more work is needed to determine how species interactions may moderate adverse climate effects, or whether there are areas at risk of being unable to adapt to changing conditions. A more holistic understanding of climatic limits at the local and regional levels will help inform more relevant climatic models and conservation strategies. Understanding the ecosystem thresholds for both hydroclimatic limits will be critical for predicting biome transitions and for providing strategies to preserve biodiversity and support communities that rely on these dryland ecosystems.

## CRediT authorship contribution statement

Arturo Sanchez-Azofeifa: Conceptualization, Methodology, Project administration, Writing – original draft, Writing – review and editing; Kayla Stan: Formal analysis, Methodology, Visualization, Writing – original draft, Writing – review and editing; Hendrik Hamann: Data curation, Formal analysis, Methodology, Writing – original draft, Writing – review and editing.

## Acknowledgements

The authors would like to acknowledge the contributions of Antonio Guzman for his work in exploratory data analysis.

## Data Availability Statement

Rasters of linear regressions of each essential climatic variable are available for download on Harvard Dataverse (https://doi.org/10.7910/DVN/QRNS6L).

## References

Abdaki, M., Sanchez-Azofeifa, A., Hamann, H.F., Ludwig, R.L. Projected Air Temperature Dynamics in a Tropical Dry Forest Under NEX-GDDP-CMIP6 Scenarios. Earth Syst Environ 2025. 10.1007/s41748-025-00715-x

Abdi, H., & Williams, L. J. 20100. Principal component analysis. Wiley interdisciplinary reviews: computational statistics, 2(4), 433–459.

Andrade, E. M., Guerreiro, M. J. S., Palácio, H. A. Q., & Campos, D. A. 2020. Ecohydrology in a Brazilian tropical dry forest: thinned vegetation impact on hydrological functions and ecosystem services. Journal of Hydrology: regional studies, 27, 100649. 10.1016/j.ejrh.2019.100649

Bojinski, S., Verstraete, M., Peterson, T. C., Richter, C., Simmons, A., & Zemp, M. 2014. The concept of essential climate variables in support of climate research, applications, and policy. Bulletin of the American Meteorological Society, 95(9), 1431–1443. 10.1175/BAMS-D-13-00047.1

Bonshoms, M., Ubeda, J., Liguori, G., Körner, P., Navarro, Á., & Cruz, R. 2022. Validation of ERA5-Land temperature and relative humidity on four Peruvian glaciers using on-glacier observations. Journal of Mountain Science, 19(7), 1849–1873. 10.1007/s11629-022-7388-4

Calvo-Rodríguez, S., Sanchez-Azofeifa, A. G., Durán, S. M., & Espírito-Santo, M. M. 2017. Assessing ecosystem services in Neotropical dry forests: a systematic review. Environmental Conservation, 44(1), 34–43. 10.1017/S0376892916000400

Castro, S. M., Sanchez-Azofeifa, G. A., & Sato, H. 2018. Effect of drought on productivity in a Costa Rican tropical dry forest. Environmental Research Letters, 13(4), 045001. 10.1088/1748-9326/aaacbc

De Marzo, T., Pflugmacher, D., Baumann, M., Lambin, E. F., Gasparri, I., & Kuemmerle, T. 2021. Characterizing forest disturbances across the Argentine Dry Chaco based on Landsat time series. International Journal of Applied Earth Observation and Geoinformation, 98, 102310. 10.1016/j.jag.2021.102310

Deb, J. C., Phinn, S., Butt, N., & McAlpine, C. A. 2017. The impact of climate change on the distribution of two threatened Dipterocarp trees. Ecology and evolution, 7(7), 2238–2248. 10.1002/ece3.2846

Dinerstein, E., Olson, D., Joshi, A., Vynne, C., Burgess, N. D., Wikramanayake, E.,… & Saleem, M. 2017. An ecoregion-based approach to protecting half the terrestrial realm. BioScience, 67(6), 534–545. 10.1093/biosci/bix014

Dommo, A., Vondou, D. A., Philippon, N., Eastman, R., Moron, V., & Aloysius, N. 2022. The ERA5’s diurnal cycle of low-level clouds over Western Central Africa during June–September: Dynamic and thermodynamic processes. Atmospheric Research, 280, 106426. 10.1016/j.atmosres.2022.106426

Dong, T., Zhu, X., Deng, R., Ma, Y., & Dong, W. 2022. Detection and attribution of extreme precipitation events over the Asian monsoon region. Weather and Climate Extremes, 38, 100497. 10.1016/j.wace.2022.100497

Doughty, C. E., Keany, J. M., Wiebe, B. C., Rey-Sanchez, C., Carter, K. R., Middleby, K. B.,… & Fisher, J. B. 2023. Tropical forests are approaching critical temperature thresholds. Nature, 621(7977), 105–111. 10.1038/s41586-023-06391-z

Funk, C., Peterson, P., Landsfeld, M., Pedreros, D., Verdin, J., Shukla, S.,… & Michaelsen, J. 2015. The climate hazards infrared precipitation with stations—a new environmental record for monitoring extremes. Scientific data, 2(1), 1–21. 10.1038/sdata.2015.66

González-M, R., Posada, J. M., Carmona, C. P., Garzón, F., Salinas, V., Idárraga-Piedrahita, Á.,… & Salgado-Negret, B. 2021. Diverging functional strategies but high sensitivity to an extreme drought in tropical dry forests. Ecology Letters, 24(3), 451–463. 10.1111/ele.13659

Guo, J., Hu, S., & Guan, Y. 2022. Regime shifts of the wet and dry seasons in the tropics under global warming. Environmental Research Letters, 17(10), 104028. 10.1088/1748-9326/ac9328

Guzmán Q., J. A., Hamann, H. F., & Sánchez-Azofeifa, G. A. 2024. Multi-decadal trends of low clouds at the Tropical Montane Cloud Forests. Ecological Indicators, 158, 111599. 10.1016/j.ecolind.2024.111599

Hartung, M., Carreño-Rocabado, G., Peña-Claros, M., & van der Sande, M. T. 2021. Tropical dry forest resilience to fire depends on fire frequency and climate. Frontiers in Forests and Global Change, 4, 755104. 10.3389/ffgc.2021.755104

Hersbach, H., Bell, B., Berrisford, P., Hirahara, S., Horányi, A., Muñoz-Sabater, J.,… & Thépaut, J. N. 2020. The ERA5 global reanalysis. Quarterly journal of the royal meteorological society, 146(730), 1999–2049. 10.1002/qj.3803

Hollunder, R., Garbin, M. L., Rubio Scarano, F., & Mariotte, P. 2022. Regional and local determinants of drought resilience in tropical forests. Ecology and Evolution, 12(5), e8943. 10.1002/ece3.8943

Hong, P., Schmid, B., De Laender, F., Eisenhauer, N., Zhang, X., Chen, H.,… & Wang, S. 2022. Biodiversity promotes ecosystem functioning despite environmental change. Ecology Letters, 25(2), 555–569. 10.1111/ele.13936

Husson, F., Josse, J., Le, S., & Mazet, J. 2024. FactoMineR: Multivariate exploratory data analysis and data mining (Version 2.11) [R package]. https://cran.r-project.org/package=FactoMineR

Jiao, T., Williams, C. A., De Kauwe, M. G., Schwalm, C. R., & Medlyn, B. E. 2021. Patterns of post-drought recovery are strongly influenced by drought duration, frequency, post-drought wetness, and bioclimatic setting. Global Change Biology, 27(19), 4630–4643. 10.1111/gcb.15788

Kanniah, K. D., Beringer, J., & Hutley, L. B. 2013. Response of savanna gross primary productivity to interannual variability in rainfall: Results of a remote sensing-based light use efficiency model. Progress in Physical Geography, 37(5), 642–663. 10.1177/0309133313490006

Kim, J. B., Kim, S. H., & Bae, D. H. 2023. The impacts of global warming on arid climates and drought features. Theoretical and Applied Climatology, 152(1), 693–708. 10.1007/s00704-022-04348-2

Knapp, A. K., Ciais, P., & Smith, M. D. 2017. Reconciling inconsistencies in precipitation–productivity relationships: implications for climate change. New Phytologist, 214(1), 41–47. 10.1111/nph.14381

Kurten, E. L., Bunyavejchewin, S., & Davies, S. J. 2018. Phenology of a dipterocarp forest with seasonal drought: Insights into the origin of general flowering. Journal of Ecology, 106(1), 126–136. 10.1111/1365-2745.12858

Lu, S., Shao, X., Freitag, M., Klein, L. J., Renwick, J., Marianno, F. J.,… & Hamann, H. F. 2016. IBM PAIRS curated big data service for accelerated geospatial data analytics and discovery. In 2016, the IEEE International Conference on Big Data (Big Data) (pp. 2672-2675). IEEE.

MacFarland, T. W., & Yates, J. M. 2016. Kruskal–Wallis H-test for one-way analysis of variance (ANOVA) by ranks. In Introduction to nonparametric statistics for the biological sciences using R (pp. 177–211). Cham: Springer International Publishing. 10.1007/978-3-319-30634-6_6

Manrique-Ascencio, A., Prieto-Torres, D. A., Villalobos, F., & Guevara, R. 2024. Climate-driven shifts in the diversity of plants in the Neotropical seasonally dry forest: Evaluating the effectiveness of protected areas. Global Change Biology, 30(4), e17282. 10.1111/gcb.17282

Mendivelso, H. A., Camarero, J. J., Gutiérrez, E., & Zuidema, P. A. 2014. Time-dependent effects of climate and drought on tree growth in a Neotropical dry forest: Short-term tolerance vs. long-term sensitivity. Agricultural and Forest Meteorology, 188, 13–23. 10.1016/j.agrformet.2013.12.010

Miles, L., Newton, A. C., DeFries, R. S., Ravilious, C., May, I., Blyth, S.,… & Gordon, J. E. 2006. A global overview of the conservation status of tropical dry forests. Journal of biogeography, 33(3), 491–505. 10.1111/j.1365-2699.2005.01424.x

Moura, M. R., do Nascimento, F. A., Paolucci, L. N., Silva, D. P., & Santos, B. A. 2023. Pervasive impacts of climate change on the woodiness and ecological generalism of dry forest plant assemblages. Journal of Ecology, 111(8), 1762–1776. 10.1111/1365-2745.14139

Nelson, H. P., Devenish-Nelson, E. S., Rusk, B. L., Geary, M., & Lawrence, A. J. 2020. A review of tropical dry forest ecosystem service research in the Caribbean–gaps and policy implications. Ecosystem services, 43, 101095. 10.1016/j.ecoser.2020.101095

O’Brien, M. J., Hector, A., Ong, R., & Philipson, C. D. 2024. Tree growth and survival are more sensitive to high rainfall than drought in an aseasonal forest in Malaysia. Communications Earth & Environment, 5(1), 179. 10.1038/s43247-024-01335-5

Olson, D. M., Dinerstein, E., Wikramanayake, E. D., Burgess, N. D., Powell, G. V., Underwood, E. C.,… & Kassem, K. R. 2001. Terrestrial Ecoregions of the World: A New Map of Life on Earth: A new global map of terrestrial ecoregions provides an innovative tool for conserving biodiversity. BioScience, 51(11), 933–938. https://doi.org/10.1641/0006-3568(2001)051[0933:TEOTWA]2.0.CO;2

Pagotto, M. A., Aragão, J. R. V., Hornink, B., Menezes, I. R. N., Tomazello-Filho, M., Lisi, C. S.,… & Tabarelli, M. 2025. Biomass production by tree species is negatively affected by decreased precipitation and chronic anthropogenic disturbance in a Caatinga dry forest. Journal of Arid Environments, 228, 105340. 10.1016/j.jaridenv.2025.105340

Park, C. E., Jeong, S. J., Joshi, M., Osborn, T. J., Ho, C. H., Piao, S.,… & Feng, S. 2018. Keeping global warming within 1.5°C constrains the the emergence of aridification. Nature Climate Change, 8(1), 70–74. 10.1038/s41558-017-0034-4

Pennington, R. T., Lehmann, C. E., & Rowland, L. M. 2018. Tropical savannas and dry forests. Current Biology, 28(9), R541–R545.

Perez, S., Hammond, M., & Lau, J. 2025. Precipitation anomalies may affect productivity resilience by shifting plant community properties. Journal of Ecology, 113(3), 542–554. 10.1111/1365-2745.14471

Poorter, L., Rozendaal, D. M., Bongers, F., Almeida, D. J. S., Álvarez, F. S., Andrade, J. L.,… & Westoby, M. 2021. Functional recovery of secondary tropical forests. Proceedings of the National Academy of Sciences, 118(49), e2003405118. 10.1073/pnas.2003405118

Portillo-Quintero, C. A., & Sánchez-Azofeifa, G. A. 2010. Extent and conservation of tropical dry forests in the Americas. Biological Conservation, 143(1), 144–155. 10.1016/j.biocon.2009.09.020

Rankine, C., Sanchez-Azofeifa, A., do Espirito-Santo, M. M., & Stan, K. 2024. Succession and seasonality of a Brazilian secondary tropical dry forest: Phenology and climate moderation. Forest Ecology and Management, 568, 122151. 10.1016/j.foreco.2024.122151

Schröder, J. M., Rodríguez, L. P. Á., & Günter, S. 2021. Research trends: Tropical dry forests: The neglected research agenda?. Forest Policy and Economics, 122, 102333. 10.1016/j.forpol.2020.102333

Siyum, Z. G. 2020. Tropical dry forest dynamics in the context of climate change: syntheses of drivers, gaps, and management perspectives. Ecological Processes, 9(1), 1–16. 10.1186/s13717-020-00229-6

Stan, K., & Sanchez-Azofeifa, A. 2019. Tropical dry forest diversity, climatic response, and resilience in a changing climate. Forests, 10(5), 443. 10.3390/f10050443

Stan, K. D., Sanchez-Azofeifa, A., Duran, S. M., Hesketh, M., Laakso, K., Portillo-Quintero, C.,… & Doetterl, S. 2021. Tropical dry forest resilience and water use efficiency: an analysis of productivity under climate change. Environmental Research Letters, 16(5), 054027. 10.1088/1748-9326/abf6f3

Stan, K. D., Sanchez-Azofeifa, A., & Hamann, H. F. 2024. Widespread degradation and limited protection of forests in global tropical dry ecosystems. Biological Conservation, 289, 110425. 10.1016/j.biocon.2023.110425

Suresh, H. S., Dattaraja, H. S., & Sukumar, R. 2010. Relationship between annual rainfall and tree mortality in a tropical dry forest: results of a 19-year study at Mudumalai, southern India. Forest Ecology and Management, 259(4), 762–769. 10.1016/j.foreco.2009.09.025

Wang, L., & Collins, S. L. 2024. The complex relationship between precipitation and productivity in drylands. Cambridge Prisms: Drylands, 1, e1. 10.1017/dry.2024.1

Walter, J. A., Atkins, J. W., & Hulshof, C. M. 2024. Climate and topography control variation in the tropical dry forest–rainforest ecotone. Ecology, 105(11), e4442. 10.1002/ecy.4442

Werden, L. K., Averill, C., Crowther, T. W., Calderón-Morales, E., Toro, L., Alvarado, J. P.,… & Powers, J. S. 2023. Belowground traits mediate tree survival during tropical dry forest restoration. Philosophical Transactions of the Royal Society B, 378(1867), 20210067. 10.1098/rstb.2021.0067

Xie, Y., Wang, X., & Silander Jr, J. A. 2015. Deciduous forest responses to temperature, precipitation, and drought imply complex climate change impacts. Proceedings of the National Academy of Sciences, 112(44), 13585–13590. 10.1073/pnas.1509991112

Xu, H., Lian, X., Slette, I. J., Yang, H., Zhang, Y., Chen, A., & Piao, S. 2022. Rising ecosystem water demand exacerbates the lengthening of tropical dry seasons. Nature Communications, 13(1), 4093. 10.1038/s41467-022-31826-y

Yang, L. Y., Yu, R., Wu, J., Zhang, Y., Kosugi, Y., Restrepo-Coupe, N.,… & Tan, Z. H. 2023. Asian tropical forests assimilating carbon under dry conditions: water stress or light benefits?. Journal of Plant Ecology, 16(3), rtac106. 10.1093/jpe/rtac106

Yu, Z., Wang, J., Liu, S., Rentch, J. S., Sun, P., & Lu, C. 2017. Global gross primary productivity and water use efficiency changes under drought stress. Environmental Research Letters, 12(1), 014016. 10.1088/1748-9326/aa5258

Zhang, Y., Gentine, P., Luo, X., Lian, X., Liu, Y., Zhou, S.,… & Keenan, T. F. 2022. Increasing sensitivity of dryland vegetation greenness to precipitation due to rising atmospheric CO2. Nature Communications, 13(1), 4875. 10.1038/s41467-022-32631-3

Zou, L., Cao, S., Zhao, A., Sanchez-Azofeifa, A. 2020. Assessing the temporal response of tropical dry forests to meteorological drought. Remote Sensing. 12, 2341. doi:10.3390/rs12142341

